# Effect of Compound Kushen Injection, a natural compound mixture, and its identified chemical components on migration and invasion of colon, brain and breast cancer cell lines

**DOI:** 10.1101/500124

**Authors:** Saeed Nourmohammadi, Thazin Nwe Aung, Jian Cui, Jinxin V. Pei, Michael Lucio De Ieso, Yuka Harata-Lee, Zhipeng Qu, David L Adelson, Andrea J Yool

## Abstract

Cancer metastasis is a major cause of death. Traditional Chinese medicines (TCM) are promising sources of new anti-metastatic agents. Compound Kushen Injection (CKI), extracted from medicinal plants, Kushen (Sophora flavescens) and Baituling (Heterosmilax chinensis), contains a mixture of alkaloids and flavonoids known to disrupt cell cycle and induce apoptosis in breast cancer (MCF7). However, effects on cancer cell migration and invasion have remained unknown. CKI, fractionated mixtures, and single identified components were tested in migration assays with colon (HT-29, SW-480, DLD-1), brain (U-87 MG, U-251 MG), and breast (MDA-MB-231) cancer cell lines. Human embryonic kidney (HEK-293) and human foreskin fibroblast (HFF) served as non-cancerous controls. Wound closure, transwell invasion, and live cell imaging assays showed that CKI reduced motility in all eight cell lines. The greatest inhibition of migration occurred in HT-29 and MDA-MB-231, and the least in HEK-293. Fractionation and reconstitution of CKI showed that combinations of compounds were required for activity. Live cell imaging confirmed CKI strongly reduced migration of HT-29 and MDA-MB-231 cells, moderately slowed brain cancer cells, and had no effect on HEK-293. CKI uniformly blocked invasiveness through extracellular matrix. Apoptosis was increased by CKI in MDA-MB-231 cells but not in non-cancerous cells. Cell viability in CKI was unaffected in all cell lines. Transcriptomic analyses of MDA-MB-231 with and without CKI indicated down-regulated expression of actin cytoskeletal and focal adhesion genes, consistent with the observed impairment of cell migration. The pharmacological complexity of CKI is important for its effective block of cancer cell migration and invasion.

## Introduction

Cancer progression results from uncontrolled migration of cells away from the primary tumor, intravasation into lymphatic or vascular circulation, invasion into secondary tissues and the formation of metastasized tumors (1, 2) which are the main cause of cancer-related deaths (3–5). Migrating cells generate a driving force for cell motility based on the extension of filipodia and lamellipodia using actin polymerization at the leading edges of the cell (6). New approaches for enhancing cancer treatment by impairing cell migration and metastasis might offer promise for curing patients with malignancies. Combinations of anticancer therapeutic regimens that not only reduce cell proliferation but also limit metastasis would be advantageous (7–9).

Herbal medicines are used in complementary and traditional medicines (10–12). Traditional Chinese medicine (TCM) relies on natural product extracts containing complex mixtures of components, suggested to deliver therapeutic benefit to individuals suffering from non-small cell lung cancer, liver, breast, and colorectal cancers (12–14). The inherent complexity of TCM suggests principle components might act in concert with adjuvant components, explaining an apparent synergy in therapeutic benefits seen from the whole extract as compared to individual compounds (15). Modulation of multiple regulatory signaling targets has been proposed as essential for the anti-proliferative, anti-migratory and anti-metastatic properties of TCMs (16, 17).

Compound Kushen Injection (CKI) has been used in combination with chemotherapies such as oxaliplatin and 5-fluorouracil in China since 1995 for the treatment of gastric, liver and non-small cell lung carcinomas (18). Composed of alkaloids, flavonoids, organic acids and saccharides (19), CKI has been reported to boost immunity, decrease inflammation, and decrease metastasis (20), for example by repressing RNA markers associated with tumor metastasis in MCF-7 cells (17, 18), and impairing migration in hepatocellular carcinoma cells (21). The challenge for defining mechanisms of action of TCMs such as CKI is to understand the differential activities of chemical components not only singly but in combination, recognizing the likely involvement of multiple gene expression and signaling pathways in the beneficial outcomes (22).

Work here is the first to show that CKI and defined chemical fractions slow cancer cell migration and invasion, and to use systems biology to identify sets of genes linked to cell migration that are regulated by CKI treatment. Differential expression of genes in the actin cytoskeleton and focal adhesion pathways supports the idea that the therapeutic activity of CKI in humans involves a serendipitous combination of effects on cancer cell properties.

## Materials and Methods

### Cell lines

MDA-MB-231, HT-29, SW-480, DLD-1, U-87 MG, U-251 MG and HEK-293 were purchased from the American Type Culture Collection (ATCC, Manassas, VA) and human foreskin fibroblast (HFF) was kindly provided by Dr. Eric Smith (Basil Hetzel Institute, The Queen Elizabeth Hospital, SA, Australia). DLD-1 cells were grown in RPMI (Roswell Park Memorial Institute) culture medium (Thermo Fisher Scientific, MA, USA) with 10 % fetal bovine serum (FBS, Thermo Fisher Scientific). All other cell lines were cultured in Dulbecco Modified Eagle Medium (DMEM, Thermo Fisher Scientific) with 10 % FBS, except for HFF (which contained 15 % FBS), at 37^0^C in 5 % CO_2_.

### CKI preparation and other chemicals

CKI (Batch No: 20170322, total alkaloid concentration of 25 mg/ml) was obtained from Zhendong Pharmaceutical Co. Ltd (Shanxi, China). High performance liquid chromatography (HPLC) fractionation and liquid chromatography-mass spectrometry (LC-MS/MS) were used to analyse single compounds and confirm their concentrations in CKI. Fractionation of CKI was done by using Shimadzu HPLC SPC-M20A photodiode-array UV-Vis detector (Japan) equipped with a C_18_ column (5 μm, 250 x 10 mm; Phenomenex, CA, USA), with methods for fractionation and concentration determinations as described previously (23) (Supplementary Figs 2 and 3). The nine compounds comprising the major fraction (MJ): oxymatrine, oxysophocarpine, n-methylcytisine, matrine, sophocarpine, trifolirhizin, adenine, and sophoridine were purchased as isolated compounds from Beina Biotechnology Institute Co., Ltd (Shanxi, China), and macrozamin was obtained from Zhendong Pharmaceutical Co. Ltd (Beijing, China). Two-fold serial dilutions from 2 mg/ml through to 0.25 mg/ml of total alkaloids in CKI, as well as equivalent dilutions of the MJ and minor (MN) fractions were used for bioassay experiments on the cancer and non-cancerous cell lines and compared with effects of vehicle control treatments; 0.25 *%* Tween 80 (Sigma-Aldrich, MO, USA) and 10 mM HEPES (Thermo Fisher Scientific) in the same cell lines.

### Circular wound closure assay

Two-dimensional (2D) cell migration was measured using circular wound closure rates (24). CKI-based and vehicle control treatments were applied in low serum DMEM, with the mitotic inhibitor 5-fluoro-2'-deoxyuridine (FUDR). Initial wound areas were imaged at 0 h (10x objective) with a Canon EOS 6D camera (Canon, Tokyo, Japan) on an Olympus inverted microscope (Olympus Corp., Tokyo, Japan); XnConvert software was used to standardize the images. NIH ImageJ software (U.S. National Institutes of Health, MD, USA) was used to quantify wound areas at 0 h and at a second timepoint (18 to 24 h), set for each cell line to ensure wounds were not completely closed. Experiments were independently repeated three times, with four to eight replicates.

### Transwell invasion assay

Invasion assays were performed in 24-well transwell inserts (6.5 mm, 8 μm pore size; Corning^®^ Transwell polycarbonate; Sigma-Aldrich). The upper surface of the filter was layered with extracellular matrix (ECM) gel from Sigma-Aldrich (at dilutions empirically optimized for each cell line; Supplementary Table 2), allowed to dry overnight, and rehydrated with 50 μl of serum-free media per insert for 1 h prior to cell seeding. Cultures were grown to 40 % confluence and then starved in medium with 2 % FBS serum for 24 h prior to seeding. Cells were detached (at ≤ 80 % confluency) and resuspended in serum-free culture media (Supplementary Table 2). Cells were then seeded in transwell inserts (total 150 μl of cell suspension per transwell, including 50 μl of rehydration medium added earlier) at appropriate number of cells per well in the presence of CKI, MN or MJ at 2 mg/ml, or vehicle control medium. The chemoattractant gradient was created with 700 μl of culture medium (containing the relevant CKI-based or vehicle treatment), with 5 %, 10 %, or 15 % FBS for HEK-293, cancerous cells, or HFF, respectively. Depending on invasion characteristics, cells from each line were incubated for an optimized time period ranging from 5 to 24 h at 37°C in 5 % CO2. Non-invasive cells were scraped from the upper surface of the filter with a cotton swab; migrated cells on the bottom surface were counted after staining with crystal violet (Sigma-Aldrich). The average number of invasive cells was calculated from three randomly selected fields (x100 magnification). Cell counts were normalized to numbers in the vehicle control treatment. Three independent experiments were carried out with two replicates.

### Cytotoxicity assay

Cell viability was measured with the Alamar Blue assay (25) according to the manufacturer's instructions (Thermo Fisher Scientific). Cells were seeded in 96-well plates in media with 2 % FBS and FUDR. After overnight incubation, treatments were applied, and cultures were incubated for 24 h. After application of 10 % Alamar Blue solution (30-90 min), fluorescence was measured by using FLUOstar Optima microplate reader. Mercuric chloride (2.5 mM) served as a positive control, inducing cytotoxic cell death. A control sample with medium and treatment agent only (no cells) was included for background color subtraction.

### Apoptosis assay

Apoptosis assays based on annexin V and propidium iodide staining were performed as described previously (17). Briefly, MDA-MB-231, HEK-293 and HFF cells were seeded in 6-well trays and treated with 2 mg/ml of CKI. After 24 h of treatment, cells were harvested, and levels of apoptotic cells were measured using the Annexin V-FITC detection kit (Thermo Fisher Scientific) according to the manufacturer's guidelines. Cells were acquired on a LSRFortessa X-20 (BD Biosciences, NJ, USA) and data were analyzed using FlowJo software (TreeStar Inc., OR, USA).

### Live cell imaging

Cells in 96 well plates were cultured to 80 % confluency, then serum-starved for 12-18 h in optimal culture media with 2 % FBS and 400 nM FUDR. Wounds were created as circular lesions in the confluent monolayers, and treatments were added as described for the circular wound closure assay above. Plates were placed in an enclosed humidified chamber at 37°C with 5 % CO_2_ for 20 h, and images were acquired at 15-min intervals with a Nikon Ti E Live Cell Microscope (Nikon, Tokyo, Japan) using Nikon NIS-Elements software. Time-lapse movies as AVI files were exported from NIS-Elements. ImageJ (U.S. National Institutes of Health) was used to convert the exported files into TIFF files and converted files were then analyzed using Fiji software (26).

### Immunofluorescence labelling and confocal microscopy

Cells were plated in μ-Plate 8 Well dishes (Ibidi, Munich, Germany), in 2 % FBS with FUDR 400 nM in optimal culture media and incubated 12-18 h at 37°C in 5 % CO_2_. The following day, 2 mg/ml of CKI, MN, MJ or vehicle control treatments were applied, and cells were incubated for 24 h. After washing with phosphate-buffered saline (PBS), cells were fixed in 4 % paraformaldehyde (at room temperature for 10-30 min), rinsed 2-3 times with PBS, and permeabilized with 200 μl of 0.1 % Triton X-100 in PBS (3-5 min at room temperature). After 2-3 washes with PBS, Phalloidin-iFluor 488 Reagent CytoPainter (ab176753; Abcam, MA, USA) at 200 μl per well was used to stain F-actin cytoskeleton (room temperature in the dark for 1-2 h) and washed again 2-3 times with PBS. Cell nuclei were labelled with 200 μl of 1:1000 Hoechst stain (cat # 861405; Sigma-Aldrich) for 5-10□min. Cells were visualized using a SP5 laser scanning confocal microscope (Leica, Germany).

### Pathway enrichment analysis of migratory genes affected by CKI

RNA-seq (RNA-sequencing) data (23) of MDA-MB-231 cells treated with CKI were used to identify cell migration genes affected by CKI. Gene Ontology (GO) enrichment analyses were carried out with R package clusterProfiler 3.8.0 (27). The following parameters for GO enrichment analysis were used: biological process at the third level; right-sided hypergeometric test; and Benjamini-Hochberg method to correct p values. Pvalue cutoff 0.05 and FDR (false discovery rate) 0.1 values were used to identify significantly over-represented GO terms. Pathways that were significantly perturbed by CKI treatment were identified with Signaling Pathway Impact Analysis (SPIA) (28), and visualized using the pathview package in R (29). KEGG (Kyoto Encyclopedia of Genes and Genomes) functional analysis was performed with R package clusterProfiler 3.8.0 (27) and OmicCircos v 1.18.0 (30). Venn diagrams were generated using an online tool (http://bioinformatics.psb.ugent.be/webtools/Venn/).

### Intracellular protein staining and quantification by flow cytometry

Cells cultured in 6-well trays were treated with CKI, MN or MJ as described above, harvested after 24 h, fixed and permeabilized using Nuclear Factor Fixation and Permeabilization Buffer Set (Biolegend, CA, USA) according to the manufacturer’s instructions. 2×10^5^ cells were labelled with rabbit anti-Cyclin D1 (CCND1) (92G2, Cell Signaling Technologies, MA USA), rabbit anti-β-actin (ACTB) (D6A8, Cell Signaling Technologies), rabbit anti-protein kinase B (AKT1, 2, 3) (Ab32505, Abcam) or rabbit IgG isotype control (Cell Signaling Technologies), and these antibodies were detected with antirabbit IgG-PE (Cell Signaling Technologies). For detection of β-catenin, rabbit anti-β-catenin (CTNNB1)-Alexa Fluor 647 and isotype control for CTNNB1 rabbit IgG-Alexa Fluor 647 (Abcam) were used. The cells were then sorted, and the data were acquired on a BD LSRFortessa X-20. Sorting parameters were set to gate and exclude small particles such as cell debris and large duplex cells. The data were analysed using FlowJo software.

### Statistical tests

Statistical analyses were carried out using GraphPad Prism 8 software (San Diego, CA, USA) with one-way ANOVA. Statistically significant results were represented as p<0.05 (*) or p<0.01 (**) p<0.001 (***), or p<0.0001 (****); ns (not significant). All data are shown as mean ± standard deviation (SD); n values for independent samples are indicated in italics above the x-axes in histogram figures, unless otherwise stated.

## Results

### Functional annotation of MDA-MB-231 transcriptome treated by CKI

Transcriptome (23) analyses were performed to identify over-represented Gene Ontology (GO) terms and Kyoto Encyclopedia of Genes and Genomes (KEGG) for all differentially expressed (DE) genes by comparing MDA-MB-231 gene expression profiles with and without CKI treatment (Fig 1 and Supplementary Fig 1). Differences in gene expression levels were used to identify migration related GO terms and pathways of interest, which were classified by functional roles via GO and KEGG over-representation analyses. Enriched GO terms connected to cell migration such as “positive regulation of locomotion”, “tissue migration “and “leucocyte migration” emerged from analyses of DE genes in CKI-treated MDA-MB-231 cells (Supplementary Fig 1 and Supplementary Data 1). Integration of DE genes associated with CKI treatment into KEGG pathways showed that some of the most over-represented pathways were “focal adhesion”, “regulation of actin cytoskeleton”, “pathways in cancer”, “TGF-β signaling pathway” and “adherens junction” (Fig 1). These results indicated that many of the genes affected by CKI treatment were involved in cell migration-related pathways.

**Fig. 1:**
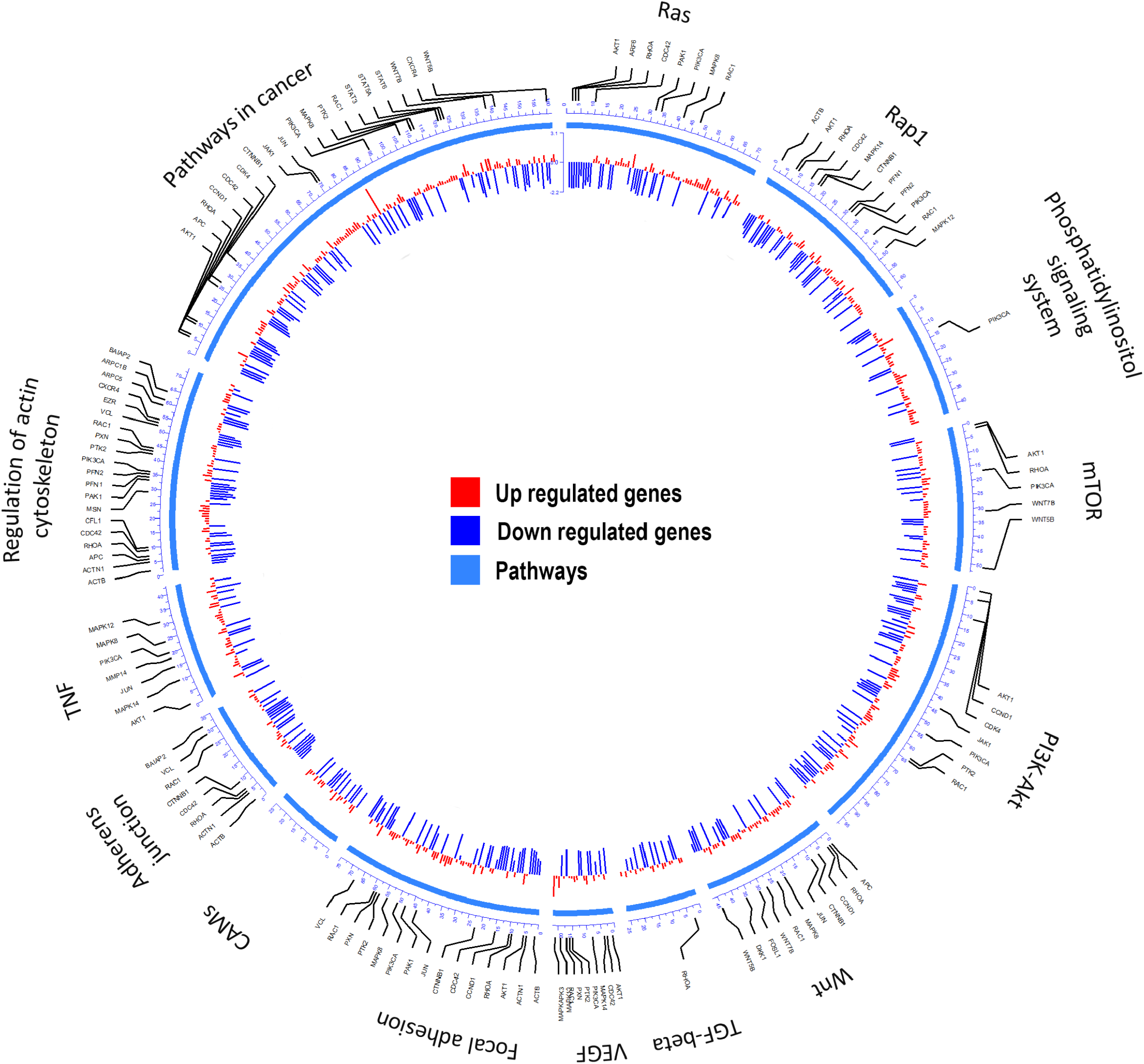
Summary of the KEGG analyses of over-represented pathways for differentially expressed genes after CKI treatment in MDA-MB-231 cells. From outer to inner, the first circle indicates the pathways; the second shows the genes involved; and the third summarizes significant changes in expression for transcript levels that were upregulated (red) or downregulated (blue) following CKI treatment. P-value cutoff for KEGG analysis was 0.05.

### Identification of CKI components

CKI was fractionated and components were selectively recombined to create MJ and MN fractions (Supplementary Fig 2) as treatments for bioassays on cultured cells. Concentrations of the nine major compounds in CKI were measured using LC-MS/MS (see Methods). The concentrations of MJ components were determined (Supplementary Figs 3 and 4) from calibration curves of nine standard compounds as previously described (23) and summarized in Supplementary Table 1. The total concentration of the nine major compounds in CKI was 9.99 mg/ml which accounted for approximately 40 % of the dry mass of CKI, and the total concentration of the nine compounds in MJ was 8.62 mg/ml, indicating that the major compounds were clearly fractionated from CKI, and that MJ and MN components were well separated.

### Impairment of cell migration by CKI and fractionated mixtures

Effects of CKI on two-dimensional cell migration were assessed using a circular wound healing assay (Fig 2A). Percent wound closure after 20 h was calculated based on the initial wound area. CKI-based treatments were tested on eight different cancer cell lines (HT-29, SW-480, DLD-1 U-87 MG, U-251 MG and MDA-MB-231) and two non-cancerous cell lines (HEK-293 and HFF), at five doses ranging from 0 to 2 mg/ml (Fig 2B). In all cell lines, net migration rates were inhibited more by CKI than by MN or MJ treatments alone, except in HEK-293 which showed low sensitivity to CKI. The retention of biological activity in the fractionated MJ and MN treatments was confirmed by demonstrating reconstituted CKI (in which MN and MJ were mixed together) was equally effective as CKI for blocking cell migration (Fig 2B). The most sensitive cell lines were breast cancer (MDA-MB-231) and colon cancer (HT-29). DLD-1 and HEK-293 cell lines were the least sensitive.

**Fig. 2:**
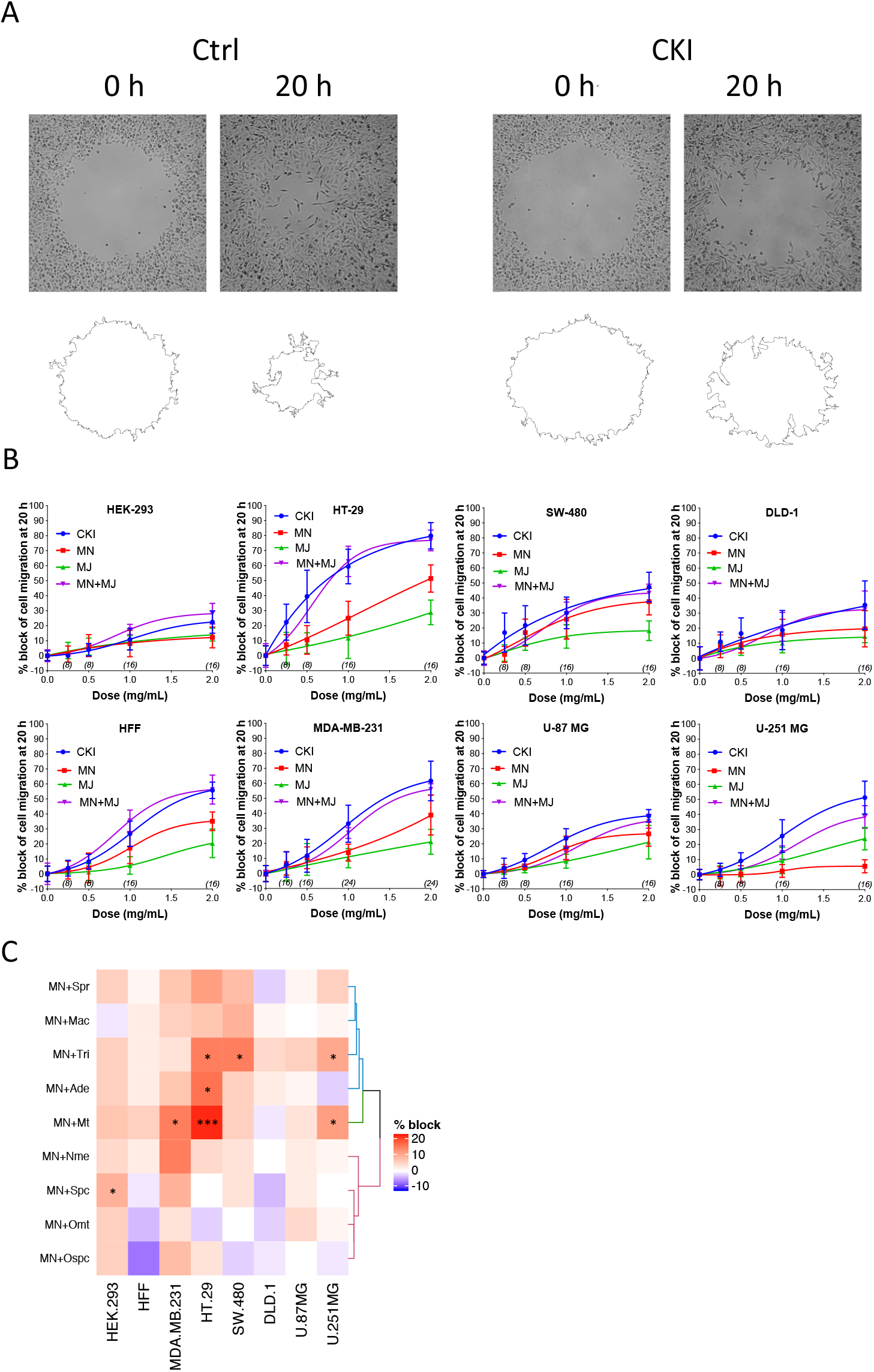
Dose-dependent inhibition of cell migration by CKI, MJ and MN fractions in eight cell lines, measured by wound closure assays. (A) Wound areas were imaged at 0 hour (initial) and after 20 hours of treatment. (B) Graphs show percent inhibition of cell migration standardized to the initial wound area, as a function of dose for treatments with CKI (blue), MJ (green), MN (red), and reconstituted CKI with major and minor fractions combined (MN+ MJ; purple). (C) Combinatorial analysis of effects on wound closure for the MN fraction tested in combination with each of nine individual major compounds of CKI, summarized as a heatmap. Data were normalized to values for percentage of migration blocked with MN alone at 0.5 mg/ml. Boxes display the net effects of added single major compounds, as no change (white), increased percentage block (red), or reduced percentage of migration blocked (blue). Statistically significant differences are shown as p < 0.05 (*), p < 0.01 (**), and p < 0.001 (***). No symbol in a box indicates the response was not significantly different from that with MN alone.

In all the CKI-sensitive cell lines, the inhibition of migration by MN alone was greater than that seen with MJ alone (Fig 2B), but neither treatment was as potent as CKI in any cell line. To determine if a single compound in MJ accounted for the enhanced inhibition seen with coapplication of MJ and MN, each of the isolated major compounds alone was added in turn to 1 mg/ml MN, at a final concentration equal to its original concentration in 2 mg/ml CKI. Wound healing assays showed certain compounds added to MN produced a significantly greater inhibition than MN alone, but the effective compounds and levels of potency differed between cell lines (Fig 2C and Supplementary Fig 5). Matrine was effective in MDA-MB-231, HT-29 and U-251 MG cell lines; sophocarpine was effective in HEK-293; trifolirhizin was effective in HT-29, SW-480 and U-251 MG; and adenine was effective in HT-29 cells, as determined by a significant decrease in migration as compared to MN alone. More than one compound in MJ appeared to contribute to the activity of CKI in blocking cell migration.

None of the CKI-based treatments induced significant cytotoxicity at 1 or 2 mg/ml, as assessed by Alamar Blue assays (Figs 3A, B, C). The lack of cytotoxicity suggested that the observed impairment of the two-dimensional cell migration was not an indirect consequence of reduced cell viability. However, CKI at 2 mg/ml substantially increased apoptosis in MDA-MB-231 and moderately increased apoptosis in HEK-293 cells, without a significant effect on HFF (Figs 3D, E, F, G). These data extend prior work which showed CKI increased apoptosis in MCF-7 breast cancer cells (Qu et al., 2016), and support the idea that multiple responses induced by CKI could contribute to its overall anti-cancer effects.

**Fig. 3:**
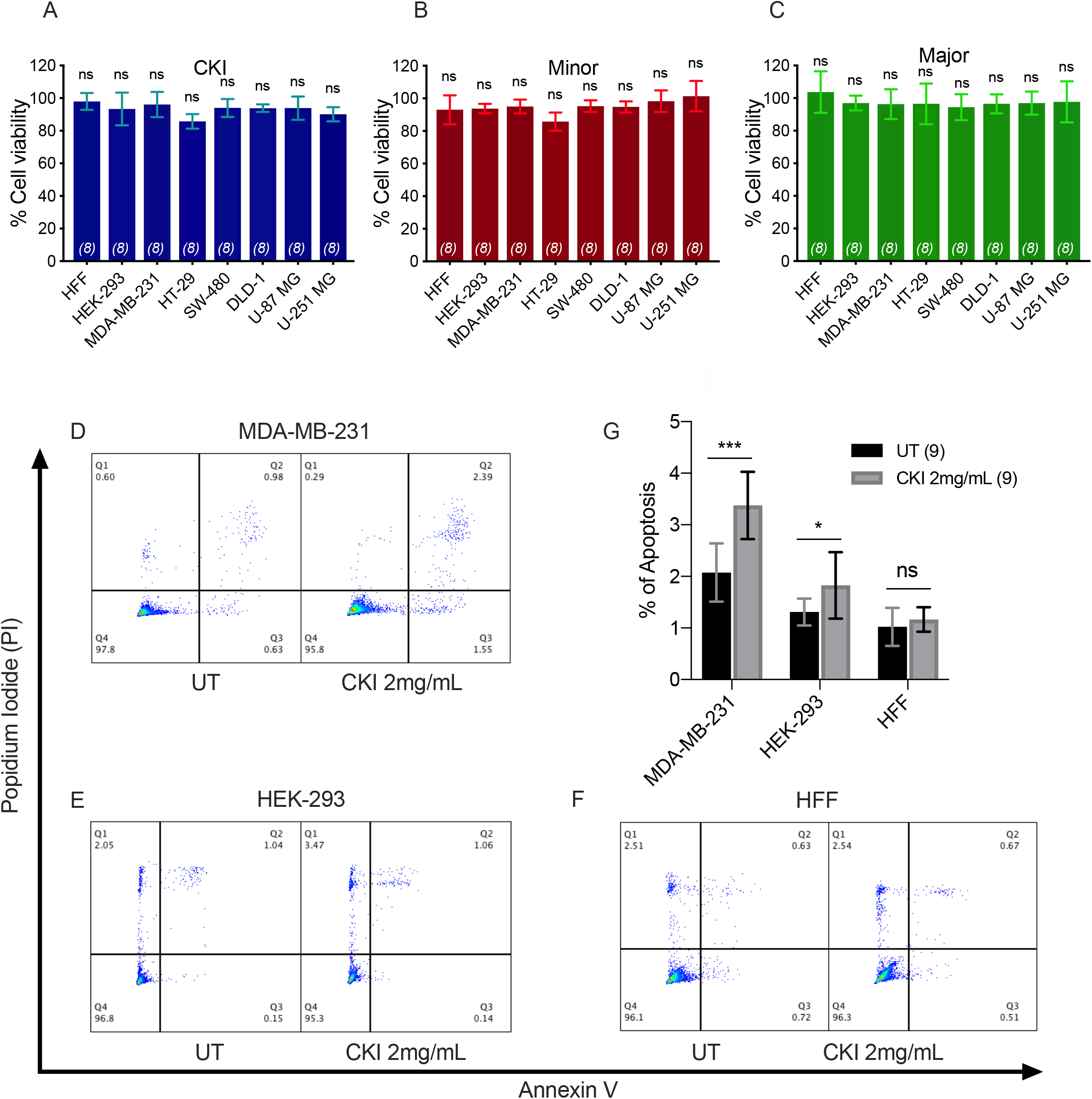
Assessment of the effects of CKI and fractions on cell viability and incidence of apoptosis. Viability was measured by Alamar Blue assay in eight cell lines treated with (A) CKI, (B) MN or (C) MJ fractions (at doses present in 2 mg/ml CKI). Cell viability responses standardized as a percentage to mean values for vehicle control were not significantly different (ns) in any condition, based on repeated experiments with 8 replicates total. Apoptosis was compared in three cell lines with and without CKI treatment, for (D) MDA-MB-231, (E) HEK-293, and (F) HFF cells, analyzed by flow cytometry (see Methods for details). Pseudo-color plots illustrate the percentages of cells in the late (quadrant Q2) and early (Q3) stages of apoptosis. (G) Histogram summarizing compiled data (mean ± SD) depicting percentage apoptosis in three cell lines with and without CKI treatment.

To quantify the effects of CKI on two-dimensional cell motility *in vitro* in more detail, trajectories of individual cells were monitored in real time using live cell imaging (Fig 4). Vehicle-treated cells were compared with those treated with 2 mg/ml CKI, or 1 mg/ml MN or MJ (doses equal to their concentration in CKI). Data were compiled for representative cells from six cancer and two non-cancerous cell lines. Positions of individual cells as a function of time over 20 h were determined by the location of the cell nucleus (Fig 4A). Distances moved per unit time interval showed a Gaussian distribution; the peak position illustrates the mean distance travelled per increment. Significant displacement of the curve to the left (representing a decreased mean distance travelled per time interval) was evident for CKI treatment in all cell lines as compared to vehicle-treated controls (Fig 4B, and Supplementary video). Significant reductions in mean distance by either MN or MJ were observed in HT-29 and SW-480 cells. Reductions in mean distance by MN but not MJ were seen in U-87 MG, MDA-MB-231 and HFF cells. U-251 MG and DLD-1 cancer cells responded only to whole CKI. The effectiveness of CKI in blocking migration depended on the simultaneous presence of multiple minor and major compounds, but the specific agents conferring anti-migration activity appeared to depend on the cell line.

**Fig. 4:**
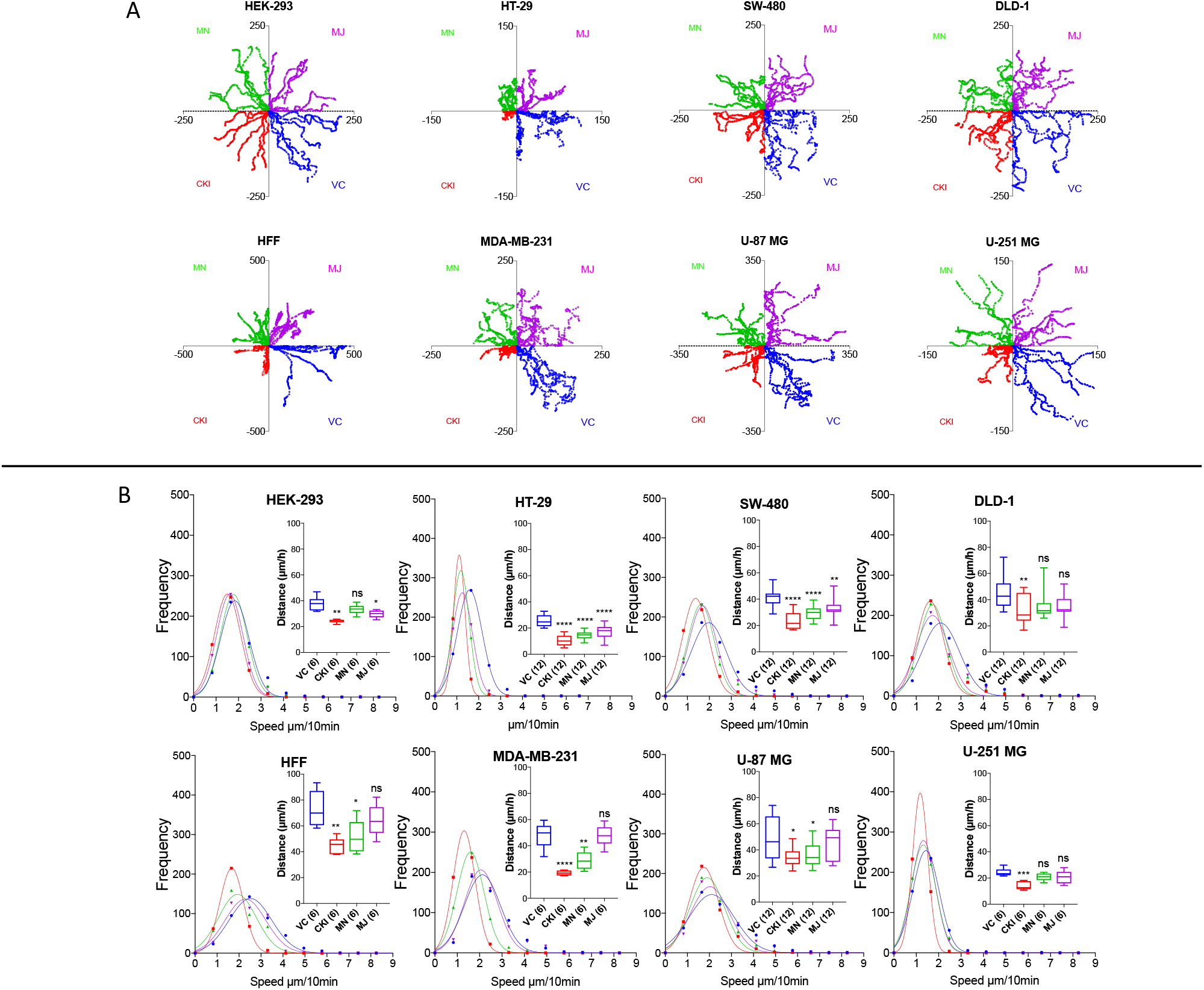
Live-cell imaging of individual cell migration trajectories, with and without the treatments in eight cell lines. Six representative cells located near wound boundaries were selected at time zero and tracked by time-lapse imaging at 10-minute intervals for 20 hours by the positions of cell nuclei. (A) Trajectory plots of individual cells, starting at the graph origin at time 0, were assigned in quadrants based on treatment groups: VC (vehicle control, blue), CKI (red), MN (green), and MJ (violet). X and Y axis values are in μm. (B) Frequency histograms of distances moved by individual cells per 10-minute interval over 20 hours. Colours indicate treatment groups, as above. Total cumulative distances moved per cell are collated in box plots (insets). Boxes enclose 50 % of values; error bars show the full range; and horizontal lines are median values. Statistically significant difference is indicated as p < 0.05 (*), p < 0.01 (**), and p < 0.001 (***).

### Impairment of invasiveness by CKI treatments in cancer and non-cancer cell lines

A transwell Boyden chamber assay with ECM-coated filters was used to measure the invasiveness of cells in response to a serum chemoattractant gradient in vehicle control, CKI, MN, and MJ treatment conditions (Fig 5). CKI treatment and the combined MN+MJ treatment (i.e., reconstituted CKI) both significantly reduced invasiveness in all cell types. Smaller but significant decreases in invasiveness were seen with MN alone in all cell types except SW-480 and were seen with MJ alone in all but SW-480 and MDA-MB-231 cell lines. These results suggested that a combination of major and minor components of CKI was required for maximal inhibition of invasiveness through extracellular matrix barriers.

**Fig. 5:**
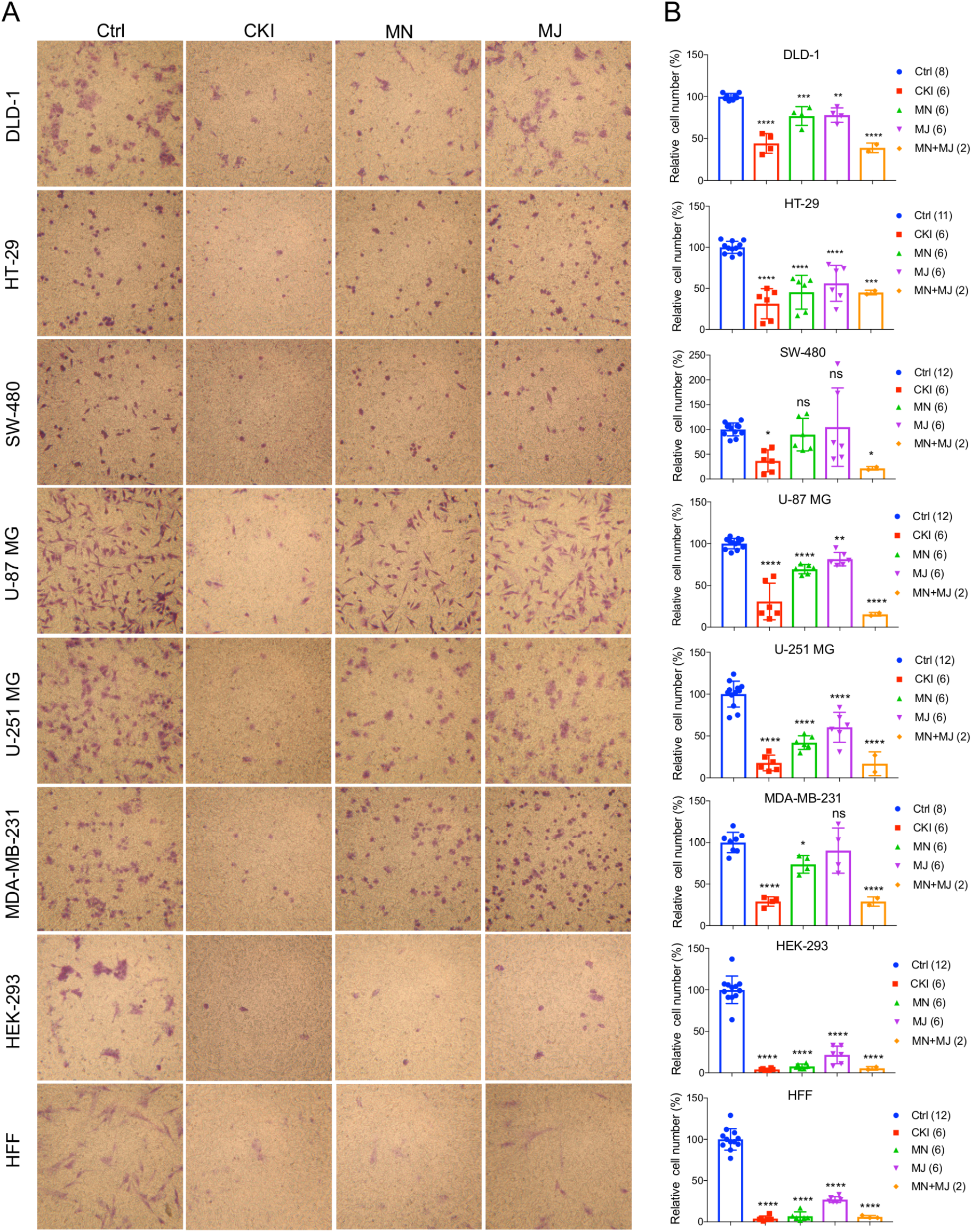
Effects of CKI and fractions on cell invasiveness in eight cell lines. Migration efficiencies of cells across an extracellular matrix barrier after treatment with CKI, minor (MN), major (MJ), or combined (MN+MJ) fractions were measured by transwell invasion assays. (A) Images show stained cells that had passed through the filter to reach the opposite side of the transwell membrane. CKI, MN and MJ were applied at doses equal to those present in 2 mg/ml CKI; Ctrl is vehicle control. (B) Compiled results are shown in histograms; n-values are in italics in corresponding figure key. Statistically significant differences compared to vehicle control are indicated as p < 0.01 (**), p < 0.001 (***), and p < 0.0001 (****); ns is not significant.

### Transcriptome analysis of MDA-MB-231 cells treated with CKI

Functional enrichment analysis was used to narrow the field of candidate mechanisms potentially associated with the anti-migratory effects of CKI. Perturbations of KEGG Pathways that were significantly over-represented in DE genes affected by CKI treatment were further analysed using SPIA (Fig 6A and Supplementary Data 2). “Melanogenesis”, “TGF-β signaling pathway”, “focal adhesion”, “regulation of actin cytoskeleton”, “ErbB signaling pathway”, and “GnRH signaling pathway” were significantly perturbed based on ‘global perturbation values’ (pG<0.05), consistent with the results of the KEGG analysis. Together, these analyses provided evidence that CKI was likely to impair cell migration by altering both adhesion and motility (Figs 6B and C). To explore this idea, genes from two strongly affected pathways (“focal adhesion” and “actin cytoskeleton”) were characterised by comparing three independent gene datasets: (i) a set containing 135 Tumor Alterations Relevant for Genomics driven Therapy (TARGET) genes; (ii) a set containing 140 migration related genes collected from published articles; and (iii) a set containing 1381 genes from KEGG pathways. We identified 14 clinically relevant DE genes, which were: *CTNNB1, CDH1, AKT1, AKT2, AKT3, CCND1, MAPK1, JAK2, APC, CDK4, RB1, PIK3CA*, and *PTEN*, all of which have been shown to affect cell migration (Fig 6D, Supplementary Table 3 and Supplementary Data 3), suggesting that the slowing of cell migration could be clinically relevant to CKI treatment outcomes.

**Fig. 6:**
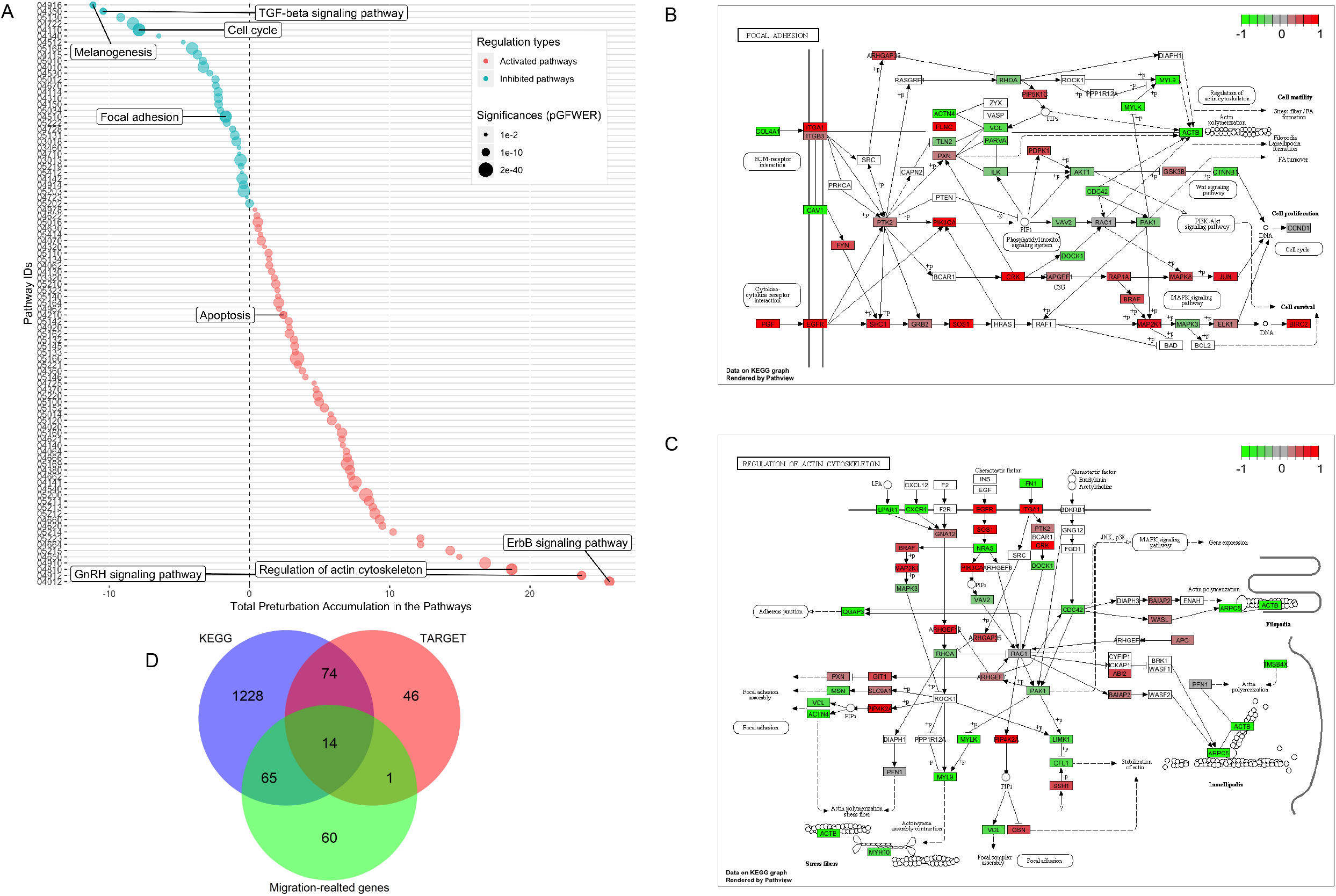
Identification of significantly perturbed pathways using SPIA analysis. (A) 101 significantly perturbed pathways in MDA-MB-231 treated by CKI 2 mg/ml were observed. Total perturbation values are shown on the x-axis, and pathway IDs are on the y-axis. The global perturbation P value (pG < 0.05) was used. (B) Significantly perturbed “focal adhesion” pathway in MDA-MB-231 treated by CKI 2 mg/ml. (C) Significantly perturbed “regulation of actin cytoskeleton” pathway in MDA-MB-231 treated by CKI 2 mg/ml. Up- and down-regulated genes are shown in red and green respectively and genes that were not affected by CKI treatment are shown in white or grey. (D) 14 “core genes” that are found across three different datasets (see Methods).

### CKI interrupts F-actin polymerization

The results of functional enrichment analysis highlighted the actin cytoskeleton and focal adhesion pathways as potential targets of CKI. As a result, we decided to examine the effect of CKI on F-actin polymerization, filopodia formation and lamellipodia extension using confocal microscopy in all eight cell lines (Fig 7). Cells in all lines treated with CKI, MN and MJ (at doses equal to those in 2 mg/ml CKI) at 24 h were smaller and lacked cellular processes such as lamellipodia as compared to vehicle control treated. Abundant lamellipodia seen in the control treatment were visibly diminished in the treatment groups. Comparing responses within the treatment groups, CKI disrupted the lamellipodia extensions more than was seen with MN or MJ alone. The impairment of F-actin polymerization was consistent with the idea that MN and MJ fractions both contribute slowing cell migration.

**Fig. 7:**
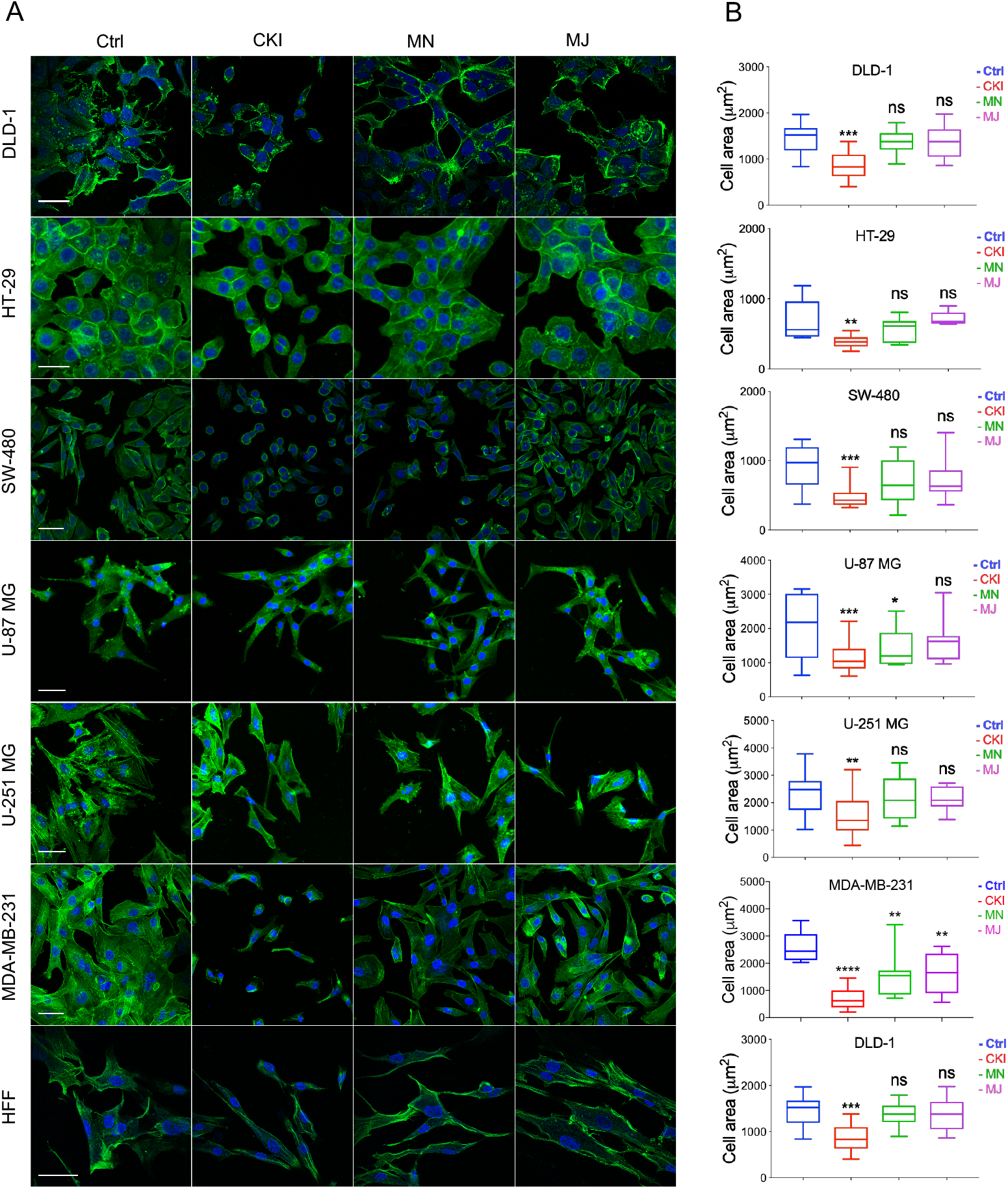
Patterns of distribution of polymerized F-actin after CKI-based treatments in eight cell lines, visualized by confocal microscopy. (A) F-actin was labelled with phalloidin (green), and nuclei with Hoechst (blue), for treatment groups (from left to right): vehicle control (Ctrl), CKI, minor and major fractions at doses present in 2 mg/ml CKI. Scale bars are 50 μm. (B) Areas (μm^2^) of F-actin staining per cell are summarized in box plots. Statistically significant results are shown as p < 0.05 (*), p < 0.01 (**), p < 0.001 (***), p < 0.0001 (****) and (#) for not significant.

### CKI and fractionated mixtures perturb the actin cytoskeleton

To validate the gene expression changes at the protein level, we performed flow cytometry analyses of four proteins; CTNNB1, AKT (1, 2, 3), CCND1, and ACTB, selected for their significant contributions in actin cytoskeleton and focal adhesion pathways in MDA-MB-231 and HEK-293 cell lines. Results shown in Fig 8 indicated that CKI, MN and MJ significantly downregulated the protein expression, confirming the gene expression data. While there was prominent down-regulation of CTNNB1 expression in MDA-MB-231 by all treatments, AKT (1, 2, 3) was downregulated by CKI and MN but not MJ treatments. In HFF cells, all treatments downregulated ACTB and CCND1 significantly; CKI and MN significantly reduced AKT (1, 2, 3) expression whereas MJ caused significant increase. In contrast, in the HFF cell line, CTNNB1 protein expression was significantly increased by CKI and MN, but significantly downregulated by MJ. In HEK cells, CKI upregulated ACTB, AKT and CCND1, whereas MN upregulated ACTB and downregulated CCND1; MJ had no significant impact on these four proteins (Fig 8). These results showed that CKI, MN and MJ significantly downregulated the four proteins in MDA-MB-231 cells, with similar although not identical results in the two non-cancer cell lines, providing additional support for the idea that CKI affects cancer cell migration by altering cytoskeletal structure.

**Fig. 8:**
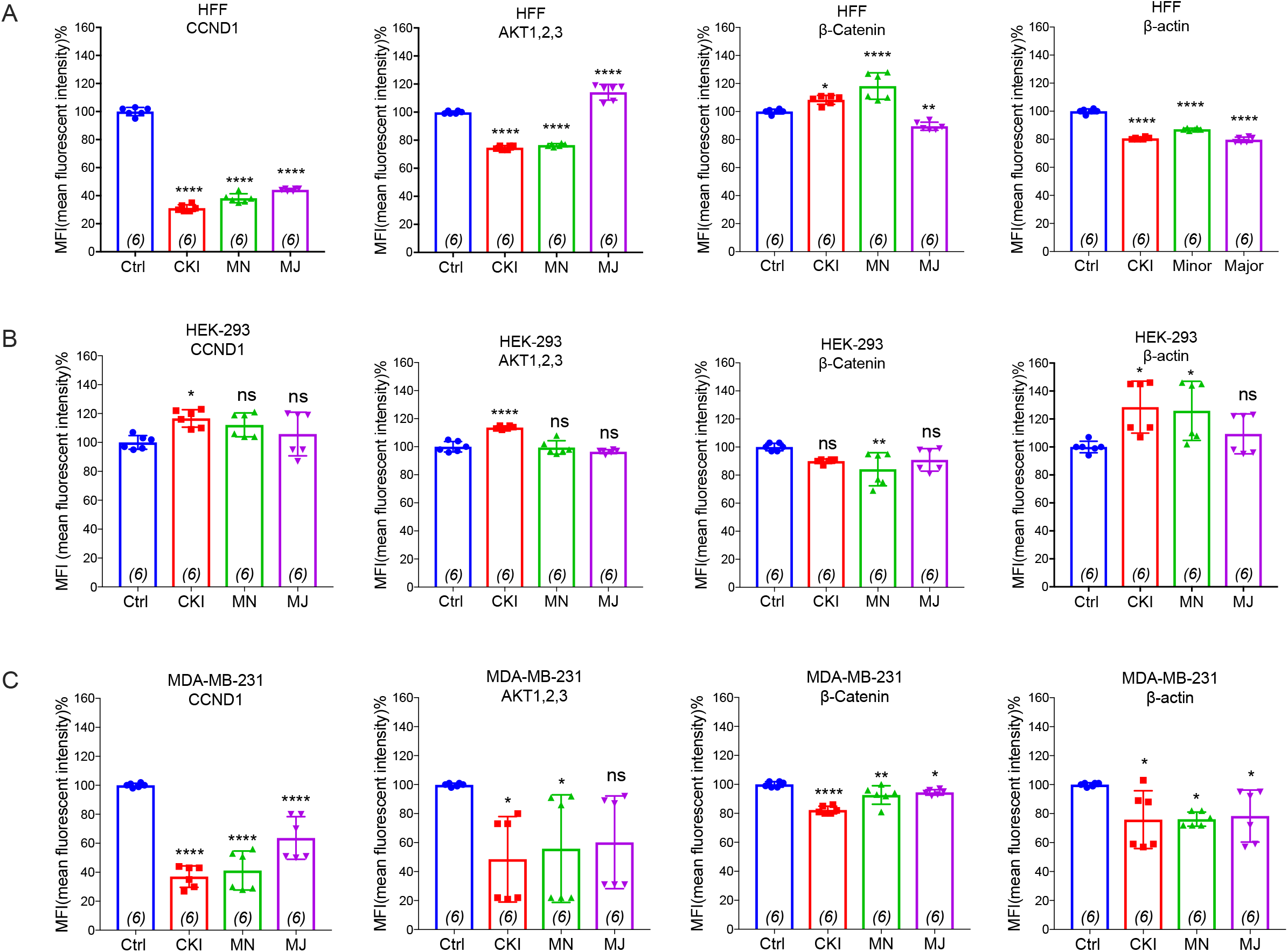
Flow cytometry analyses of four proteins: CTNNB1, AKT (1, 2, 3), CCND1, and ACTB compared in three cell lines with and without CKI-based treatments. The levels of protein expression for four genes predicted by transcriptomic analysis to be significantly down-regulated were evaluated by FACS in three cell lines (A) MDA-MB-231, (B) HEK-293, and (C) HFF. Statistically significant differences compared to vehicle controls were analysed using one-way ANOVA and post hoc tests and are shown as p < 0.05 (*), p < 0.01 (**), p < 0.001 (***), p < 0.0001 (****), and not significant (ns).

## Discussion

Natural compounds with anti-cancer activities have been suggested to include potential antimetastatic agents (17, 21, 31, 32). Clinical chemotherapeutics for cancers are targeted mainly to killing rapidly dividing cells (33), but cause serious side effects including immunosuppression, hair loss and infertility, without eliminating risks of secondary neoplasms (34). Non-toxic treatments to reduce metastasis as adjuncts to primary cancer therapies are greatly needed. CKI increased apoptotic activity in breast cancer MCF-7 and hepatocellular carcinoma HCC cell lines (17, 21). Results here showed that CKI also significantly slowed cell migration and invasion, an outcome consistent with its beneficial clinical effects. The complexity of TCM presents a challenge for identifying single active compounds. By separating components of CKI into major and minor groups, and by adding novel screens for migration and invasion into the analyses of CKI biological activity, we have shown that co-application of multiple compounds was far more effective in blocking cell migration than single agents alone, and that different compounds (serendipitously combined in CKI) are required for activity across diverse cell types.

Components of CKI have both anti-proliferation and anti-migration activities which differ between cell lines, suggesting that refinement of the TCM composition could enable customized management for different cancer types. A subset of the major compounds in CKI have been investigated previously. For example, oxymatrine impaired angiogenesis in mouse breast cancer *in vitro* and *in vivo*, by altering NF-ĸB pathway and VEGF signaling *(35).* Matrine inhibited migration and proliferation of mouse lung adenocarcinoma *in vitro* and slowed xenograft growth *in vivo*, by reducing expression of a calcium-dependent chloride channel shown to be upregulated in multiple cancer types (36, 37). Consistent with these findings, we found matrine added to the CKI minor fraction further impaired cancer cell migration (Fig 2C); however, in contrast, addition of oxymatrine to the CKI minor fraction did not affect the control of migration in any of the cell lines we tested. Oxysophocarpine slowed metastasis of oral squamous cell carcinoma *in vitro* and *in vivo*, by altering transcription factor Nrf2 and stress protein signaling pathways (38). In contrast, our results showed oxysophocarpine partially reversed the inhibition of migration observed with the CKI minor fraction. Work here showed that the major components trifolirhizin and adenine, not previously characterized, when added to CKI minor fraction further slowed cell migration, suggesting these agents merit further study as potential therapeutics, singly and in combination. Not all components of CKI enhance its anti-cancer activities; for example, depletion of three compounds (oxymatrine, oxysophocarpine and macrozamin) increased the anti-proliferative activity of CKI (23).

Rational analysis of the differential effects of agents in a TCM mixture can benefit from quantitative systems biology approaches. Transcriptomics has proven valuable for identifying altered patterns of global gene expression in response to complex agents such as CKI (18). Our pathway analysis of the transcriptome of CKI-treated MDA-MB-231 cells revealed a strong association with genes linked to “TGF-β signaling”, “focal adhesion”, “GnRH signaling”, and “regulation of actin cytoskeleton”. These pathways are linked with migratory phenotype, actin polymerization and lamellipodia protrusion (39) (40) (41). Focal adhesion sites are points of contact with extracellular matrix, anchoring actin filaments via protein complexes with transmembrane integrin receptors (42). Matrine has been reported to disrupt actin filament organization in DLD-1 colorectal adenocarcinoma cells (43). Work here is the first to show that treatment with whole CKI significantly perturbed “focal adhesion” and “regulation of actin cytoskeleton” pathways, and to confirm predictions of the transcriptomic results by showing CKI reduced lamellipodial abundance, length of extension, and area of F-actin polymerization.

Two striking outcomes of our pathway analyses were the significant negative perturbation of the cytokine TGF-β (transforming growth factor beta) and positive perturbation of GnRH (Gonadotropin-releasing hormone) signaling pathways by CKI treatment. Multifunctional TGF-β promotes the process of epithelial-mesenchymal transition, which facilitates cell migration, invasion and metastasis (44, 45). Reduced TGF-β signaling would be consistent with our findings of impaired invasion and migration with CKI treatment. Conversely, GnRH activity has been associated with attenuating migration of DU145 human prostatic carcinoma cells by remodelling the actin cytoskeleton (46). Increased GnRH signaling would be consistent with beneficial effects of CKI treatment. Further work is needed to fully understand the mechanisms of action of CKI on cell migration and invasion, the signaling pathways involved, the synergistic effects of combined therapeutic agents, and the translational potential by extending these analyses to metastatic cancer cells *in vivo*.

## Conclusion

The primary outcome of this study is the demonstration that cancer cell migration and invasion rates are significantly reduced by CKI, suggesting that therapeutic activity of CKI in human cancer patients may arise in part from downregulation of a panel of key molecular targets necessary for adhesion and motility in metastasis. The secondary outcome of this study is that multiple compounds in CKI, acting together, are responsible for this effect.

## Acknowledgements

The authors acknowledge Dr. Eric Smith for providing HFF cell line and Dr. Mohamad Kourghi, and Pak Hin Chow for valuable discussions. We also thank Dr. Agatha Labrinidis and Dr. Jane Sibbons for the facilities of Adelaide University Microscopy and technical assistance.

## Author contributions

S.N, T.N.A, D.L.A and A.J.Y designed the study, analysed the data and wrote the manuscript. S.N and T.N.A conducted the experiments and J.C, J.V.P, M.L.D, Y.H-L and Z.Q assisted with the experiments and analysis.

---

**Supplementary Fig.1:** Functional classification of genes by GO over-representation analyses. Over-represented GO terms (Biological Process, BP=3) for differentially expressed (DE) genes were identified from MDA-MB-231 cells treated by CKI. Upregulated and downregulated genes contained in each term were shown in red and green respectively. GO terms shown above the blue line were significant terms related to migration.

**Supplementary Fig. 2**: HPLC profiles of the components present in (A) CKI, (B) MJ and (C) MN fractions. Samples (50 μl at 1 mg/ml) were run through a C_18_ semi-preparative column. Numbers indicate the nine major compounds; 1: macrozamin, 2: adenine, 3: n-methylcytisine, 4: sophoridine, 5: matrine, 6: sophocarpine, 7: oxysophocarpine, 8: oxymatrine, and 9: trifolirhizin.

**Supplementary Fig. 3**: (A) Total ion chromatogram (TIC) for CKI in 1 in 100 dilution from 25 mg/ml of stock concentration. Single peaks were extracted based on the molecular mass. (B) cytisine (spike in control), (C) macrozamin, (D) adenine, (E) n-methylcytisine, (F) sophoridine and matrine (similar molecular mass with different retention time) (G) oxysophocarpine, (H) oxymatrine, (I) sophocarpine and (J) trifolirhizin.

**Supplementary Fig. 4**: (A) Total ion chromatogram (TIC) for MJ in 1 in 100 dilution from 25 mg/ml of stock concentration. Single peaks were extracted based on the molecular mass. (B) adenine, (C) cytisine (spike in control), (D) macrozamin, (E) n-methylcytisine, (F) sophoridine and matrine (similar molecular mass with different retention time) (G) oxysophocarpine, (H) oxymatrine, (I) sophocarpine and (J) trifolirhizin.

**Supplementary Fig. 5**: Combinatorial analysis of the effects of MN with each of the nine major individual compounds, analyzed in eight cell lines with wound closure assays. Data were normalized to results with 0.5 mg/ml minor (MN) alone. Significantly increased or decreased percent block of migration resulting from the addition of major compounds is shown as p < 0.05 (*), p < 0.01 (**), p < 0.001 (***) and not significant (ns). Data are mean ± SD.

**Supplementary Video**: Live-cell imaging of the migration blocking effect of CKI in MDA-MB-231 cells in the wound closure migration assay. Videos show cell motility and wound closure rate in CKI at 2 mg/ml was reduced as compared to untreated control. Images were captured at 10-minute intervals for 20 hours.

**Supplementary Data1**: Significantly over-represented functional GO terms, as determined by GO analysis of the transcriptome from CKI treated MDA-MB-231 cells. (P < 0.05).

**Supplementary Data 2**: Significantly perturbed pathways, as determined by SPIA analysis of the transcriptome from CKI treated MDA-MB-231 cells. (pG < 0.05).

**Supplementary Data 3**: Matching of genes in two strongly affected pathways (“focal adhesion” and “actin cytoskeleton”) against three independent gene datasets containing: (i) a set TARGET gene and (ii) migration related genes from published articles. 14 core DE genes from three datasets including *CTNNB1, CDH1, AKT1, AKT2, AKT3, CCND1, MAPK1, JAK2, APC, CDK4, RB1, PIK3CA*, and *PTEN*, are known to have effects on cell migration.

**Supplementary Table 1**: Concentration of 9 major compounds in CKI (Batch No:20151139) and MJ. Total alkaloid content in CKI (Batch No:20151139) = 25 mg/ml based on manufacturer’s assay. Regression line for the calculation of compounds has previously been described (Aung et al., 2018).

**Supplementary Table 2**: Concentrations of Matrigel and number of cells used for each cell line in transwell invasion assay.

**Supplementary Table 3**: Fourteen clinically relevant DE genes from three independent gene datasets (see methods).

## Abbreviations

CKI: Compound Kushen Injection
MJ: Major
MN: Minor
TCM: Traditional Chinese Medicine
HFF: Human Foreskin Fibroblast
HPLC: High Performance Liquid Chromatography
GO: Gene Ontology
FUDR: 5-fluoro-2′-deoxyuridine
RNA-seq: RNA-sequencing
KEGG: Kyoto Encyclopedia of Genes and Genomes
SPIA: Signalling Pathway Impact Analysis
ACTB: β-actin
AKT1, 2, 3: Protein kinase B
CTNNB1: β-catenin
CCND1: Cyclin D1
TARGET: Tumor Alterations Relevant for Genomics driven Therapy

