## Supplementary figures and images for "Effect of Compound Kushen Injection, a natural compound mixture, and its identified chemical components on migration and invasion of colon, brain and breast cancer cell lines"

### Supplementary Figure 1

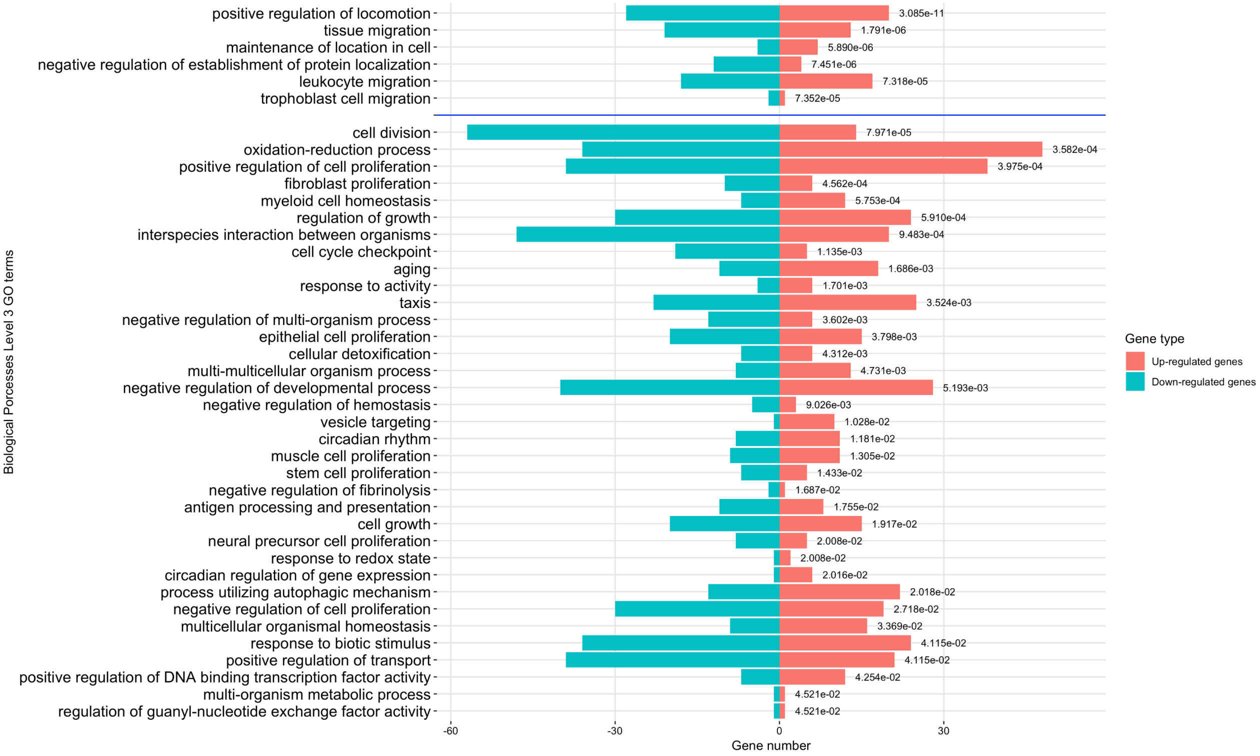

### Supplementary Figure 2

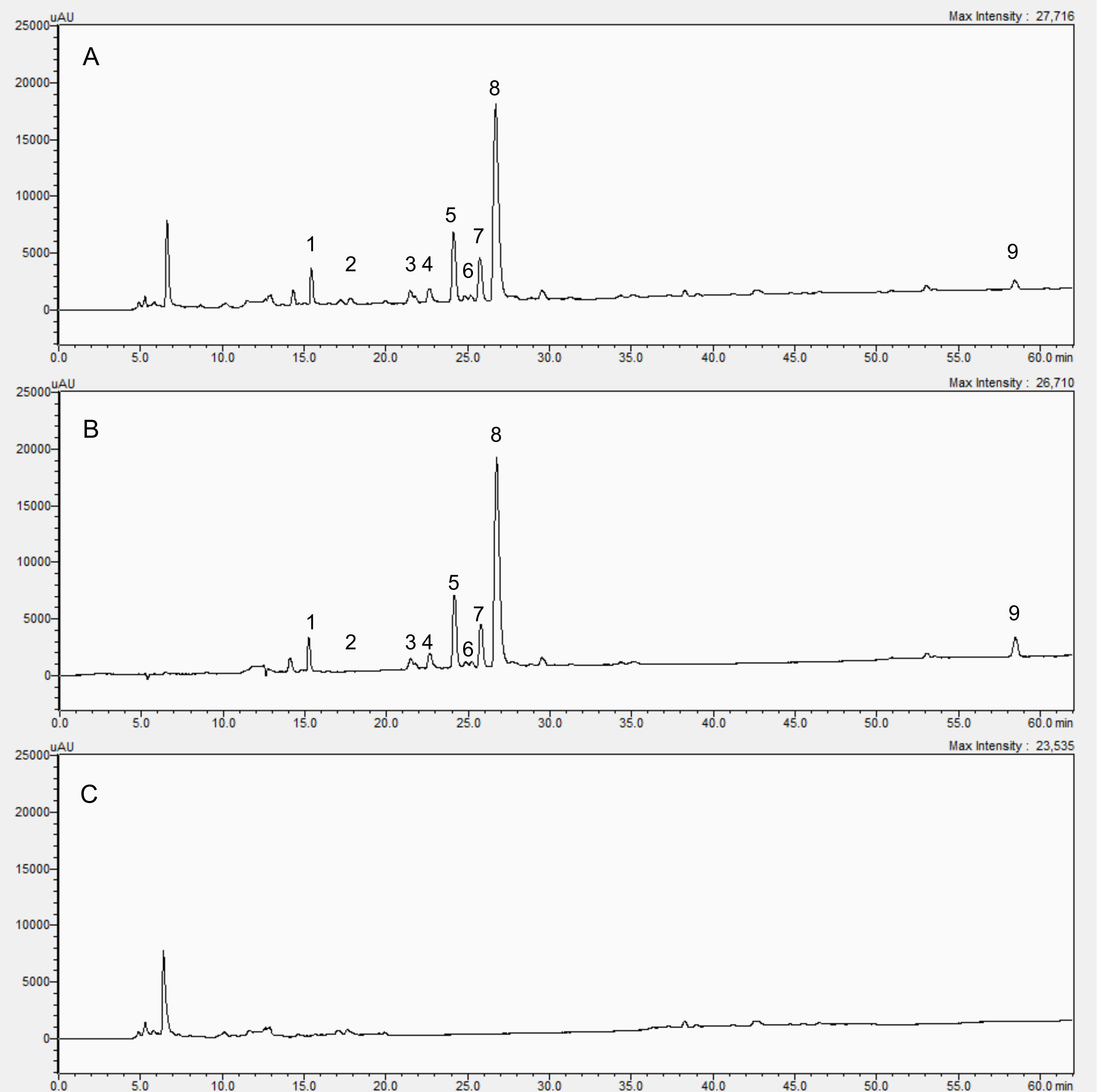

### Supplementary Figure 3

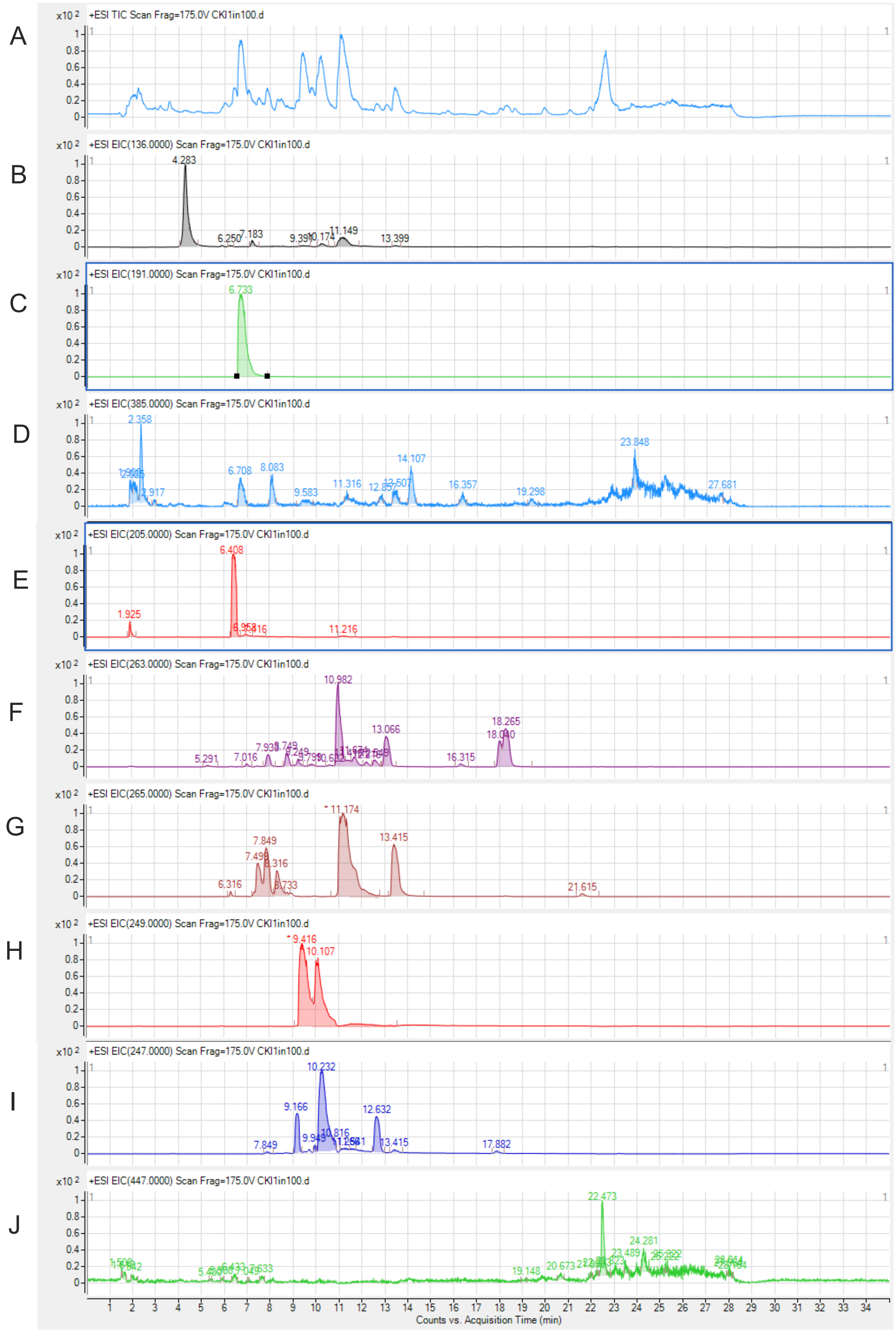

### Supplementary Figure 4

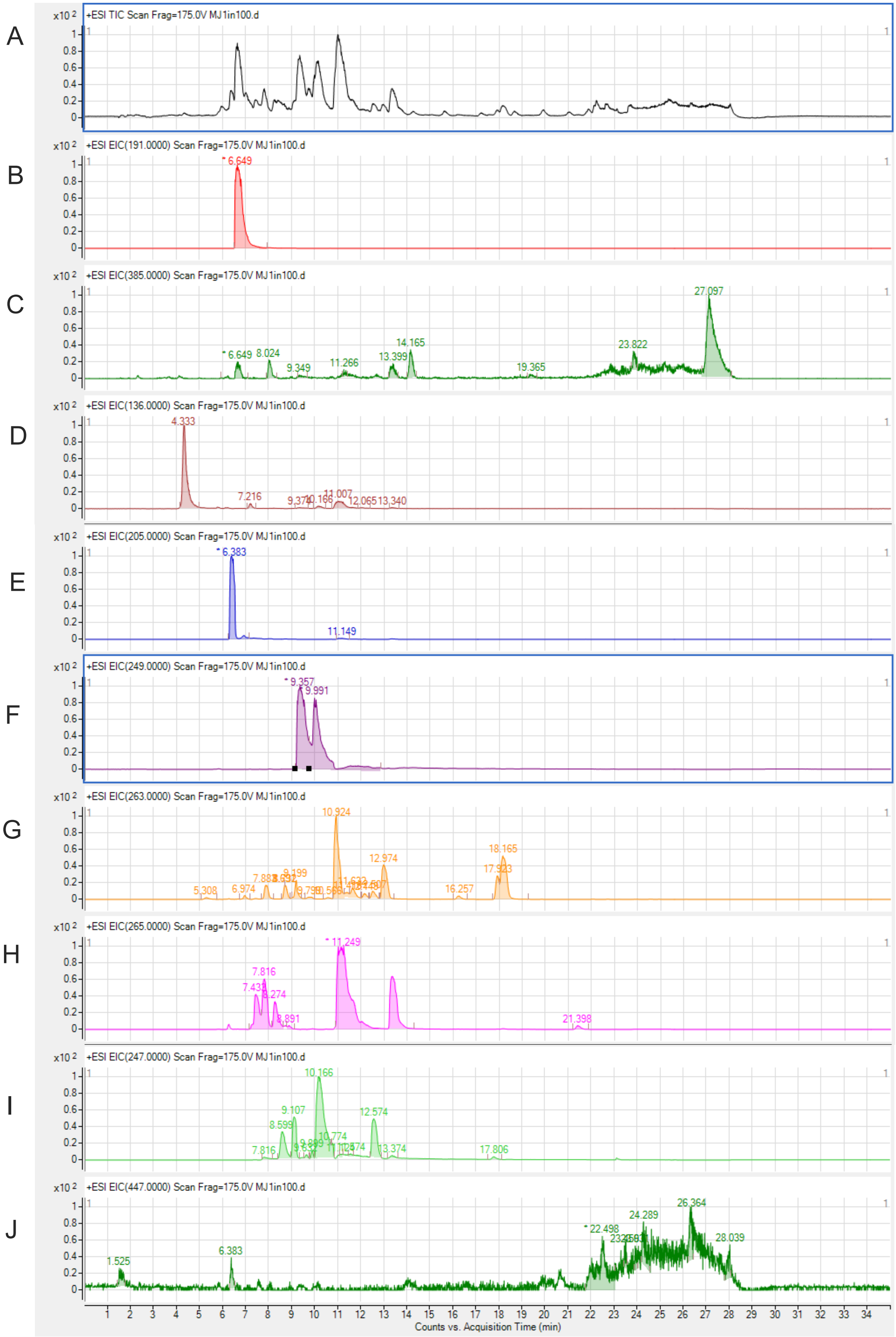

### Supplementary Figure 5

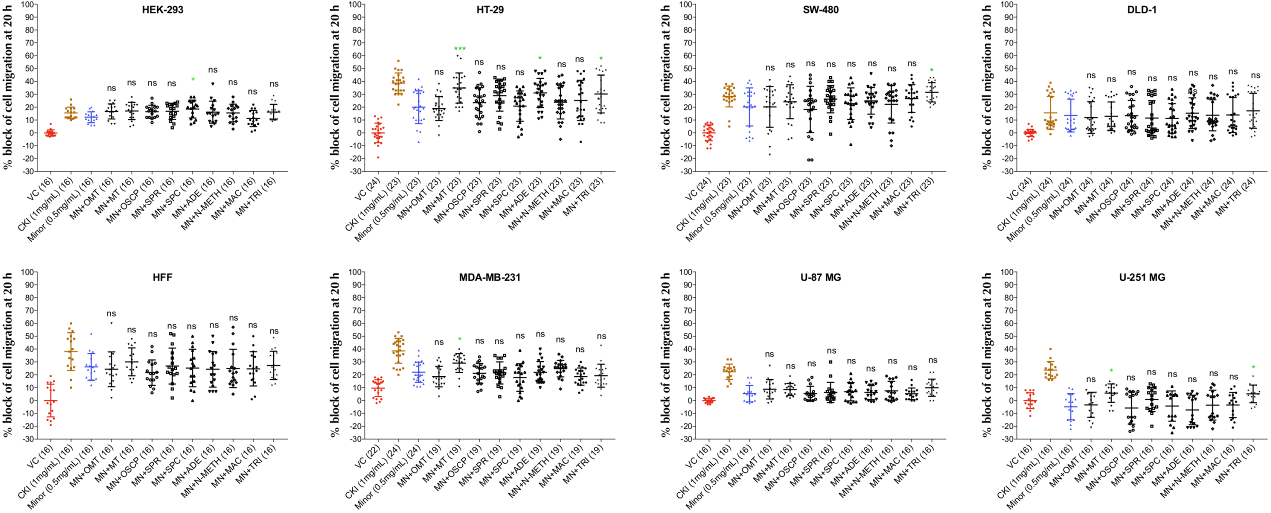
