## Supplementary Table 1 for "Effect of Compound Kushen Injection, a natural compound mixture, and its identified chemical components on migration and invasion of colon, brain and breast cancer cell lines"

**Supplementary Table 1:** Concentration of 9 major compounds in CKI (Batch No:20151139) and MJ.

| Mixtures | Compounds | concentration (mg/mL) | %Contribution |
| --- | --- | --- | --- |
| CKI | Macrozamin | 1.1 ± 0.07 | 4.4 |
|  | Adenine | 0.04 ± 0.03 | 1.6 |
|  | N-methylcytisine | 0.2 ± 0.04 | 0.8 |
|  | Sophoridine | 0.3 ± 0.19 | 1.2 |
|  | Matrine | 1.7 ± 0.17 | 6.8 |
|  | Sophocarpine | 0.5 ± 0.03 | 2 |
|  | Oxysophocarpine | 1.2 ± 0.18 | 4.8 |
|  | Oxymatrine | 5.6 ± 0.66 | 22.4 |
|  | Trifolirhizin | 0.12 ± 0.01 | 0.4 |
|  | **Total** | **11.1** | **44.4** |
| MJ | Macrozamin | 1.2 ± 0.17 | 4.8 |
|  | Adenine | 0.2 ± 0.01 | 0.8 |
|  | N-methylcytisine | 0.2 ± 0.04 | 0.8 |
|  | Sophoridine | 0.4 ± 0.12 | 1.4 |
|  | Matrine | 1.5 ± 0.6 | 6 |
|  | Sophocarpine | 0.3 ± 0.003 | 1.4 |
|  | Oxysophocarpine | 1.1 ± 0.14 | 4.3 |
|  | Oxymatrine | 5.9 ± 0.7 | 23.6 |
|  | Trifolirhizin | 0.05 ± 0.01 | 0.2 |
|  | **Total** | **10.8** | **43.3** |

*Total alkaloid content in CKI (Batch No:20151139) = 25 mg/ml based on manufacturer’s assay. Regression line for the calculation of compounds have been previously described (Aung et.al).
