## Supplementary Table 2 for "Effect of Compound Kushen Injection, a natural compound mixture, and its identified chemical components on migration and invasion of colon, brain and breast cancer cell lines"

| Cell line | Number of cells/mL | Concentration of matrigel (Stock: 10mg/mL) | Matrigel loading Volume | Incubation time (hours) |
| --- | --- | --- | --- | --- |
| MDA-MB-231 | 1X 10^6^ | 1 in 40 | 40 µL | 5 |
| U-87 | 5X 10^5^ | 1 in 30 | 40 µL | 4-5 |
| U-251 | 5X 10^5^ | 1 in 40 | 40 µL | 4-5 |
| DLD-1 | 1X 10^6^ | 1 in 100 | 40 µL | 24 |
| SW-480 | 1.5X 10^6^ | 1 in 200 | 40 µL | 24 |
| HT-29 | 2.5X 10^6^ | 1 in 400 | 40 µL | 24 |
| HEK-293 | 5X 10^5^ | 1 in 40 | 40 µL | 24 |
| HFF | 5X 10^5^ | 1 in 150 | 40 µL | 24 |

Supplementary Table 2: Concentration of Matrigel and number of cells used for each cell line in transwell invasion assay.
