## Supplementary Table 3 for "Effect of Compound Kushen Injection, a natural compound mixture, and its identified chemical components on migration and invasion of colon, brain and breast cancer cell lines"

Supplementary Table 3: Fourteen clinically relevant DE genes from three independent gene datasets.

| Gene ID | Gene Name | Full Name |
| --- | --- | --- |
| 324 | APC | APC, WNT signaling pathway regulator [ Homo sapiens (human)] |
| 595 | CCND1 | cyclin D1 [ Homo sapiens (human)] |
| 999 | CDH1 | cadherin 1 [ Homo sapiens (human)] |
| 10000 | AKT3 | AKT serine/threonine kinase 3 [ Homo sapiens (human)] |
| 1499 | CTNNB1 | catenin beta 1 [ Homo sapiens (human)] |
| 1019 | CDK4 | cyclin dependent kinase 4 [ Homo sapiens (human)] |
| 5925 | RB1 | RB transcriptional corepressor 1 [ Homo sapiens (human)] |
| 207 | AKT1 | AKT serine/threonine kinase 1 [Homo sapiens (human)] |
| 5290 | PIK3CA | phosphatidylinositol-4,5-bisphosphate 3-kinase catalytic subunit alpha [ Homo sapiens (human)] |
| 208 | AKT2 | AKT serine/threonine kinase 2 [ Homo sapiens (human)] |
| 5728 | PTEN | phosphatase and tensin homolog [ Homo sapiens (human)] |
| 5594 | MAPK1 | mitogen-activated protein kinase 1 [ Homo sapiens (human)] |
| 3717 | JAK2 | Janus kinase 2 [ Homo sapiens (human)] |
| 2099 | ESR1 | Estrogen receptor 1 [ Homo sapiens (human)] |
